# Chromosome-level genome assembly and annotation of two lineages of the ant *Cataglyphis hispanica*: stepping stones towards genomic studies of hybridogenesis and thermal adaptation in desert ants

**DOI:** 10.1101/2022.01.07.475286

**Authors:** Hugo Darras, Natalia De Souza Araujo, Lyam Baudry, Nadège Guiglielmoni, Pedro Lorite, Martial Marbouty, Fernando Rodriguez, Irina Arkhipova, Romain Koszul, Jean-François Flot, Serge Aron

## Abstract

*Cataglyphis* are thermophilic ants that forage during the day when temperatures are highest and sometimes close to their critical thermal limit. Several *Cataglyphis* species have evolved unusual reproductive systems such as facultative queen parthenogenesis or social hybridogenesis, which have not yet been investigated in detail at the molecular level. We generated high-quality genome assemblies for two hybridogenetic lineages of the Iberian ant *Cataglyphis hispanica* using long-read Nanopore sequencing and exploited chromosome conformation capture (3C) sequencing to assemble contigs into 26 and 27 chromosomes, respectively. Further karyotype analyses confirm this difference in chromosome numbers between lineages; however, they also suggest it may not be fixed among lineages. We obtained transcriptomic data to assist gene annotation and built custom repeat libraries for each of the two assemblies. Comparative analyses with 19 other published ant genomes were also conducted. These new genomic resources pave the way for exploring the genetic mechanisms underlying the remarkable thermal adaptation and the molecular mechanisms associated with transitions between different genetic systems characteristic of the ant genus *Cataglyphis*.

## Introduction

Ants of the genus *Cataglyphis* inhabit arid regions throughout the Old World, including inhospitable deserts such as the Sahara (Boulay *et al*. 2017; Lenoir *et al*. 1990). Their foraging activities are strictly diurnal, with most species being active during the hottest hours of the day (Cerda *et al*. 1998; Wehner *et al*. 1992). Some *Cataglyphis* species even forage at temperatures close to their critical thermal limits (Cerda *et al*. 1998). For instance, workers of the silver ant *Cataglyphis bombycina* have been observed to forage when ground temperatures exceed 60°C (Wehner *et al*. 1992), which supposedly provides a competitive advantage against lizard predators who avoid such harsh conditions. The high thermal tolerance seen in *Cataglyphis* species relies on a range of behavioral, morphological, physiological and molecular adaptations, such as exploitation of thermal refuges, elongated legs, high speed of movement and intense recruitment of heat-shock chaperone proteins (Aron and Wehner 2021; Gehring and Wehner 1995; Perez and Aron 2020; Perez *et al*. 2021; Pfeffer *et al*. 2019; Sommer and Wehner 2012; Willot *et al*. 2017).

In addition to their impressive heat tolerance, *Cataglyphis* ants are prominent social insect models because of their amazing diversity of reproductive traits: the number of queens per colony, the mating frequencies, the dispersal strategies and the modes of production of different castes all vary greatly among species (Aron *et al*. 2016a, 2016b; Peeters and Aron 2017). Unusual reproductive systems in which conditional use of sex produces different female castes have evolved repeatedly in different *Cataglyphis* groups. Under these systems, non-reproductive workers are sexually generated, while reproductive queens are asexually produced by thelytokous parthenogenesis – a strategy that increases the transmission rate of the genes of queens to their reproductive female offspring while maintaining genetic diversity in the worker force (Kuhn *et al*. 2020; Pearcy *et al*. 2004). Males arise from arrhenotokous parthenogenesis, as is usually the case in Hymenoptera. In several species, the conditional use of sex evolved into a unique reproductive system, named clonal social hybridogenesis, whereby male and female sexuals are produced by parthenogenesis while workers are produced exclusively from interbreeding between two sympatric, yet non-recombining genetic lineages (Darras *et al*. 2014; Eyer *et al*. 2013; Kuhn *et al*. 2020; Leniaud *et al*. 2012).

The unique characteristics of *Cataglyphis* make this ant genus an interesting model to investigate the genetic mechanisms underlying thermal adaptation and the evolution of alternative reproductive strategies. To date, only one incomplete assembly of the genome of *Cataglyphis niger*, a species characterized by classical haplodiploid reproduction, is available for genomic analyses (Yahav and Privman, 2019). To fill this gap, we combined Oxford Nanopore long reads, Illumina short reads and chromosome conformation capture (3C) sequencing (Flot *et al*. 2015; Lieberman-Aiden *et al*. 2009; Marie-Nelly *et al*. 2014) to generate high-quality chromosome-scale genome assemblies of two lineages of the Iberian ant *Cataglyphis hispanica* (Figure 1). We also annotated and compared the repeats and gene sets of this species with those of other ant genera.

**Figure 1:**
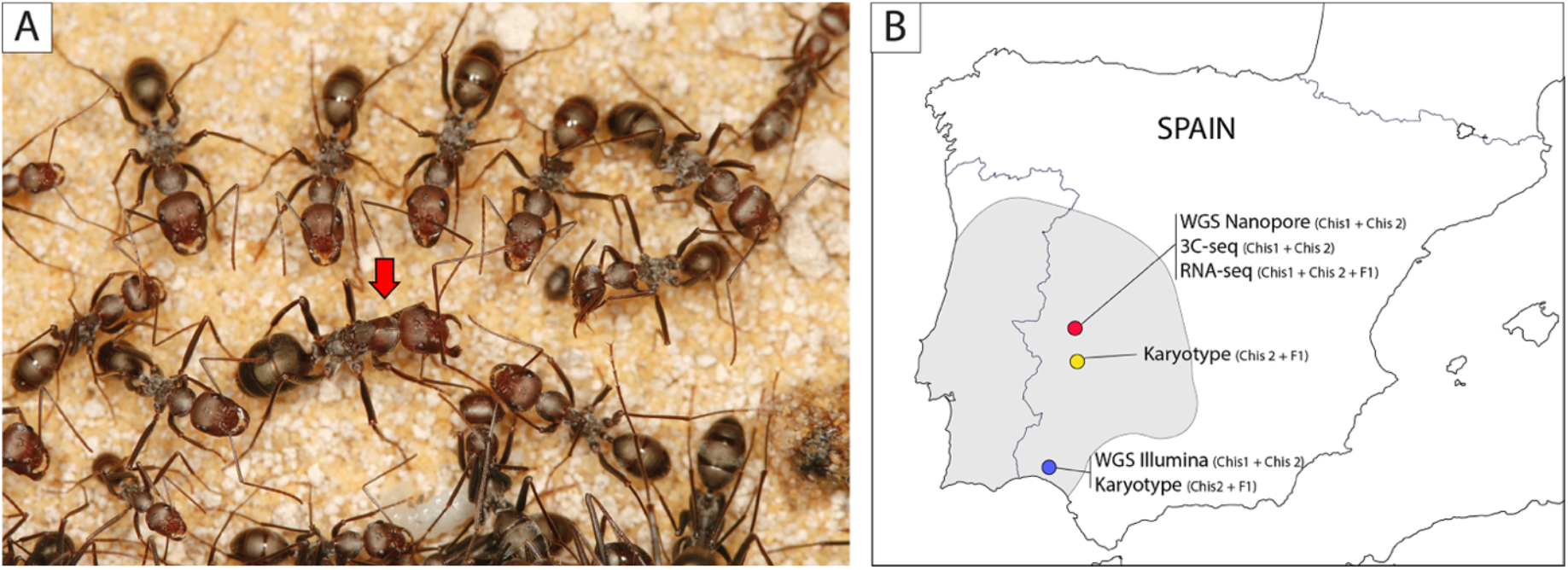
The ant *Cataglyphis hispanica*. (A) A queen of *C. hispanica* (red arrow) surrounded by workers. (B) Sampled sites in southwest Spain. The two interdependent lineages of the species, Chis1 and Chis2, were collected in Caceres (red), Merida (yellow) and Bonares (blue). For each lineage, a male from Bonares was used for whole-genome short-read sequencing (WGS Illumina) and queens from Caceres were used for both long-read sequencing (WGS Nanopore) and chromosome conformation capture sequencing (3C-seq). Karyotypes of two Chis2 males and three hybrid (F1) workers were obtained from Merida and Bonares. To assist gene annotation, transcriptomes (RNA-seq) were generated from Chis1 and Chis2 individuals from Caceres. The complete range of the species *C. hispanica* is shown in grey.

## Results and discussion

### Genome assemblies

*Cataglyphis hispanica* inhabits the most arid habitats of the Iberian Peninsula. Two sympatric hybridogenetic lineages (Chis1 and Chis2) co-occur as a complementary pair across the distribution range of the species (Leniaud *et al*. 2012; Darras *et al*. 2014). Queens of each lineage mate with males from the other lineage and produce non-reproductive workers by sexual reproduction. By contrast, male and female reproductive individuals are produced clonally through arrhenotokous and thelytokous parthenogenesis, respectively. As a result, all workers in the colonies are inter-lineage hybrids, but the two reproductive lineages do not mix genetically.

The genomes of the two hybridogenetic lineages were assembled independently (see Figure S1 for a schematic representation of the assembly pipeline). For each of the Chis1 and Chis2 lineages, we generated respectively 5.7 and 5.1 Gbp of Nanopore reads from a pool of sister clonal queens (for *de novo* long-read assemblies); 32.2 and 34.2 Gbp of PE 2 × 100 bp Illumina reads with insert sizes ranging from 170 bp to 800 bp from a single male (for short read error correction/polishing); and 8.7 and 7.0 Gbp of 3C-seq PE 2 × 66 bp (after demultiplexing) Illumina reads from a single queen (for scaffolding). The long-read assembler Flye (Kolmogorov *et al*. 2019) generated assemblies consisting of a few hundreds of contigs (439 and 929, respectively). The contigs were scaffolded using the 3C data (Marie-Nelly *et al*. 2014; Baudry *et al*. 2020): 99.7% of the Chis1 assembly was scaffolded into 26 chromosome-scale (> 2.4 Mb in length) scaffolds (Figure 2A), while 98.8 % of the Chis2 assembly was scaffolded into 27 chromosome-scale scaffolds (Figure 2B). These chromosome-scale scaffolds were labeled by decreasing size. The remaining 0.3 – 1.2% unscaffolded sequences were all relatively small (at most 34 kb for Chis1 and 119 kb for Chis2). The overall sizes of the two scaffolded assemblies were 206 Mb and 209 Mb, respectively. Assembly completeness, as estimated using BUSCO scores (Manni *et al*. 2021), was very high: among the 5,991 highly conserved single-copy genes of the Hymenoptera odb10 database, 96.8% (Chis1) and 96.1% (Chis2) were complete in each assembly. In addition, only 0.5-0.4% of the BUSCO genes appeared duplicated for both assemblies, suggesting that our assemblies did not contain much uncollapsed haplotypes, if any. In line with these results, KAT analyses based on the Illumina reads of each lineage showed a single peak of k-mer multiplicity, which were almost all represented exactly once in the assemblies as expected for high-quality genomes (Figure S2); k-mer completeness was estimated as 98.86% for Chis1 and 98.45%for Chis2 (Mapleson *et al*. 2016). For each assembly, a region with no large-scale synteny pattern was assembled at the extremity of one scaffold (the first 5.4 Mb of scaffold #9 of Chis1 and the first 3.1 Mb of scaffold #7 of Chis2). Each of these regions consisted of a collection of small contigs (mostly in the 2-10 Kb range) with 2 to 5 times higher average coverage compared to other genomic regions. These sequences exhibited microsynteny with the extremities of other large scaffolds (Figure 2 and S3) suggesting that they correspond to repeated sequences that were improperly assembled into fragmented contigs.

**Figure 2:**
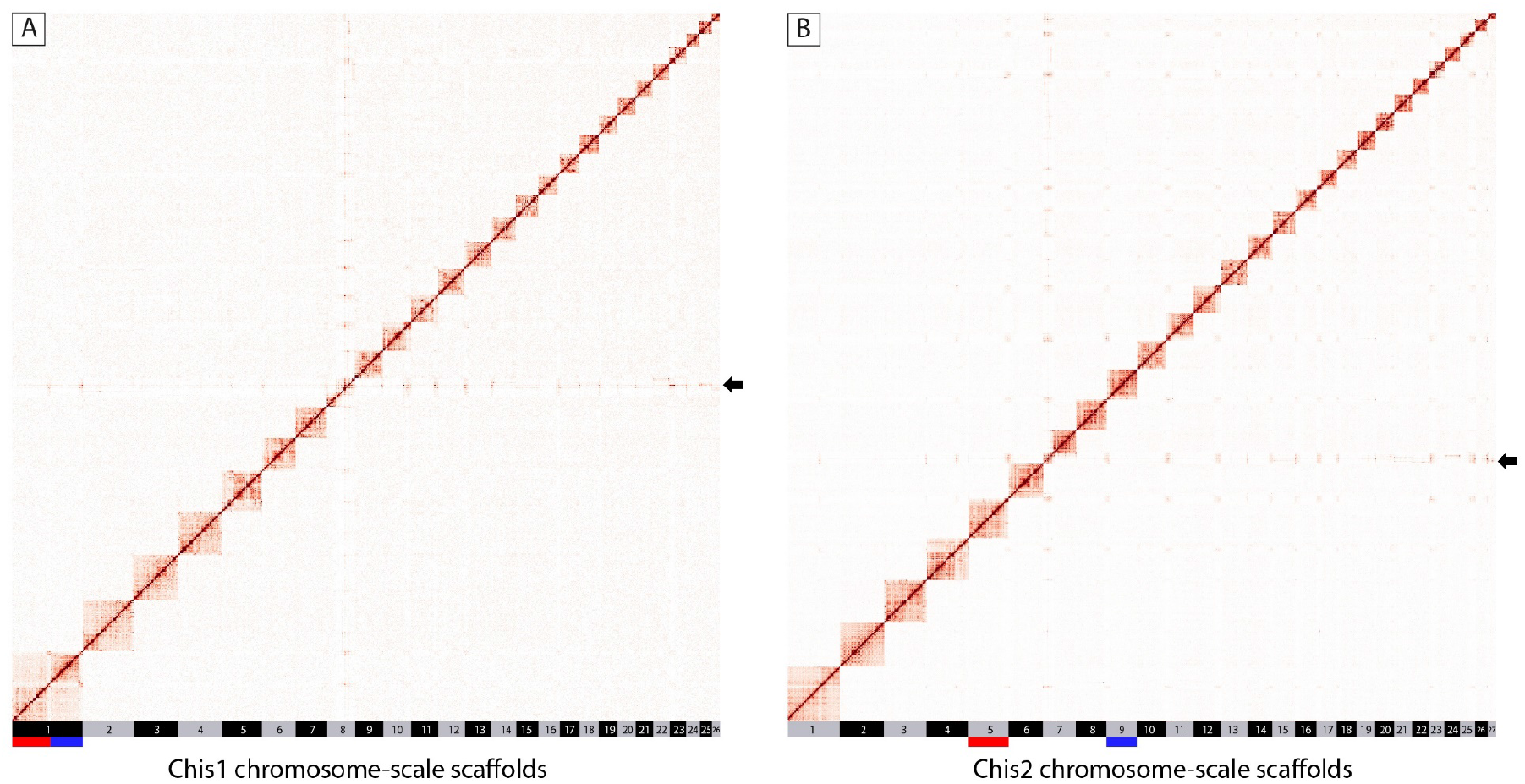
Assembly of the *Cataglyphis hispanica* Chis1 (A) and Chis2 (B) genomes into chromosomes. Hi-C interaction map revealing the presence of 26 and 27 linkage groups. The color scale represents the interaction frequencies. The positions of the rearranged chromosome are indicated, and the arrows show the assembly artefact found in each genome (see main text). The longest chromosome of Chis1 is split in two chromosomes in Chis2 (scaffolds 5 and 9, shown with red and blue colors).

Comparison of the Chis1 and Chis2 assemblies revealed that 25 of the chromosome-scale scaffolds had a one-to-one homolog in each of the two lineages. In addition, and by contrast, the largest scaffold of Chis1 (#1) was split into two chromosome-scale scaffolds (# 5 and #9) in the Chis2 assembly (Figure S3). The 3C contact maps of both lineages showed that these scaffolds (Chis1 #1 and Chis2 #5, #9) correspond to well-individualized 3D features, thereby ruling out a scaffolding error (Figure 2). These observations support that a centric fusion or fission (Robertsonian translocation) took place in one of the two lineages studied. Robertsonian translocations are the main mechanism of karyotype evolution in many animal groups, including ants (Lorite and Palomeque, 2010) and can promote speciation through the suppression of genetic recombination in the vicinity of rearranged centromeric regions or the reduction of fertility in karyotypic hybrids (Davisson and Akeson, 1993; Faria and Navarro, 2010). Intrachromosomal rearrangements between the two lineages, consisting in large translocations and inversions, were also observed for 6 of the 25 large orthologous scaffolds (Figure S3), but these could not be confirmed independently with the current data.

### Karyotyping

The numbers of chromosomes inferred for the Chis1 and Chis2 assemblies (n=26 and 27, respectively) are within the range observed in karyotypes of *Cataglyphis bicolor* (n=26), *Cataglyphis iberica* (n=26) and *Cataglyphis setipes* (n=26), as well as other Formicine species of the genera *Formica* (n=26-27), *Iberoformica* (n=26) and *Polyergus* (n=27) (Hauschteck-Jungen and Jungen, 1983; Imai *et al*. 1984; Lorite and Palomeque 2010). To determine whether the two lineages of *C. hispanica* are fixed for different chromosomal arrangements, we inspected metaphase chromosome slides from male and worker pupae from different populations (Figure 1B). In ants, as in other social Hymenoptera, males are haploid (*n*) whereas workers are diploid females (2*n*). Two males of the Chis2 lineage from Merida and Bonares were analyzed (Figure 3A and S4A-D). Both male karyotypes carried 27 chromosomes as was inferred with 3C data for the Chis2 lineage from the Caceres population. The precise morphology of the chromosomes could not be determined due to their small size (Figure 3). No male or queen pupa of the Chis1 lineage could be obtained for karyotyping. Instead, we indirectly inferred the karyotype variation in the Chis1 lineage using worker samples. Workers of *C. hispanica* are first generation hybrids and would, therefore, be expected to carry odd chromosome numbers (i.e. 2n=26+27=53) if the two lineages were fixed for different karyotypes. Workers from Bonares (N=2 from different colonies) and Merida (N=1) were analyzed. The two workers from Bonares carried 2n=54 chromosomes (Figure 3C and S4E) suggesting that the parental lineages carry the same number of chromosomes in this population. By contrast, the worker from Merida carried 53 chromosomes consistent with expectations based on genome assemblies (Figure 3E and S5). If our assumptions are correct, these results indicate that the number of chromosomes in the Chis1 lineage may vary in different populations from n=26 to n=27. The chromosomal polymorphism observed between our Chis1 and Chis2 genome assemblies is therefore unlikely to be linked to the long-term maintenance of the two lineages.

**Figure 3:**
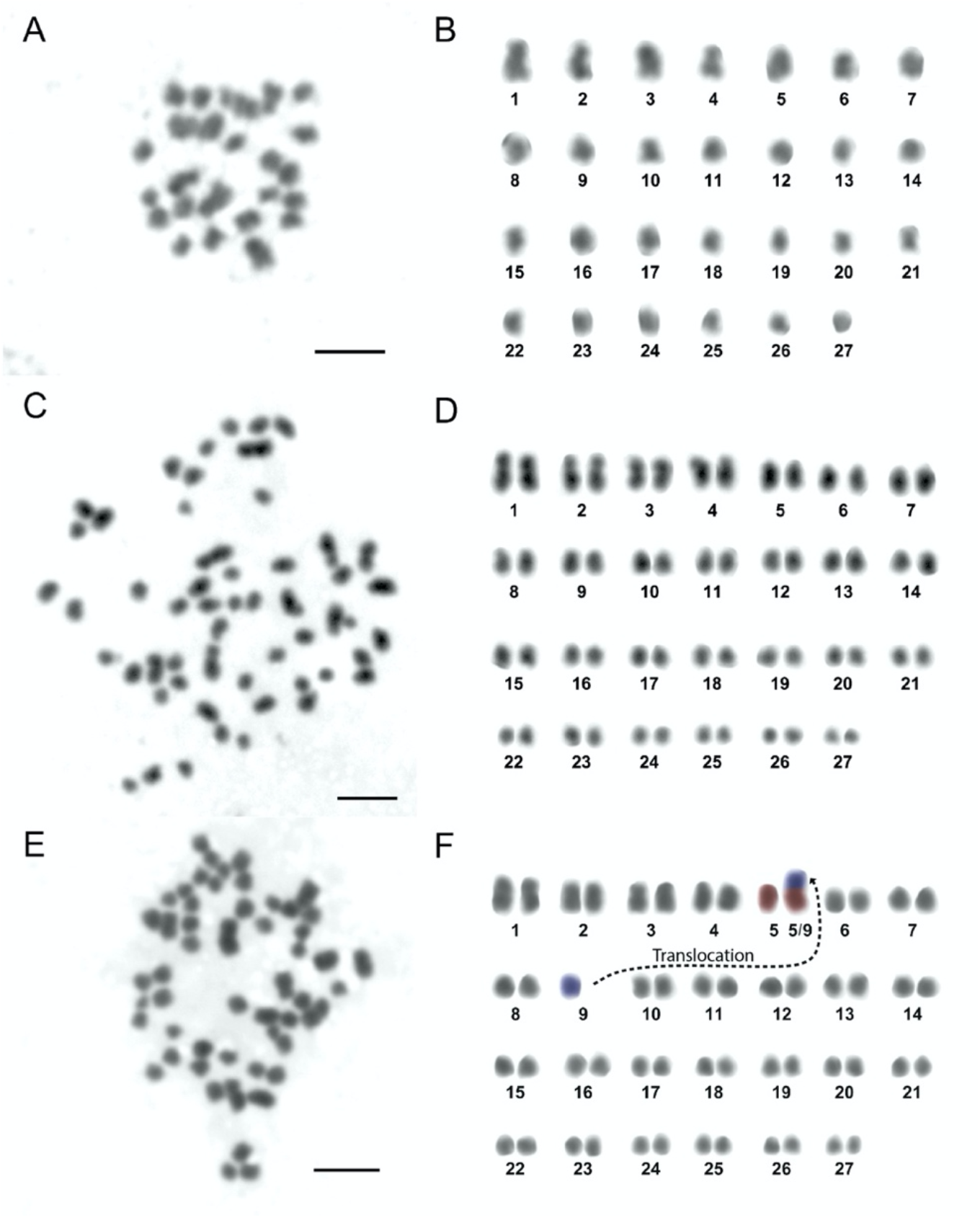
Karyotype analyses of *Cataglyphis hispanica*. (A,C and E) Metaphase chromosome slides of one haploid Chis2 male from Merida (A) and two F1 hybrid workers from Bonares (C) and Merida (E). (B,D and F) Corresponding karyotypes showing that the haploid chromosome number varies across populations. The male and the first worker display n=27 and 2n=27+27 chromosomes, respectively. By contrast, the second worker carries an unbalanced karyotype with 2n=26+27 chromosomes. The Robertsonian translocation hypothesized to be responsible for this polymorphism is indicated with dashed lines, and red and blue colors. The bar in all the images is 2 μm.

### Gene annotation

We annotated the genome of the Chis2 lineage (see Figure S1 for a schematic drawing describing the genome annotation pipeline). *Ab initio* gene prediction using AUGUSTUS and homology-based predictions using GenomeThreader (Gremme *et al*. 2005) identified 16,993 and 8,234 gene models, respectively. A total of 40,969 models (including isoforms) were also predicted by the PASA/Transdecoder (Haas *et al*. 2003) pipeline using direct evidence from 13 Gbp of Illumina RNA-seq data. The three annotation sets were validated and combined into a single annotation of 16,146 non-overlapping models using EvidenceModeler (Haas *et al*. 2008). Among these, 11,101 gene models showed significant similarity to proteins predicted in other ant species (blastp against 18 ant proteomes from the RefSeq collection) and 10,543 had functional information inferred through sequence orthology with the eggnog v5.0 database, which covers more than five thousands organisms (Huerta-Cepas *et al*. 2017, 2019). We filtered out all gene models non validated by at least one of these databases to obtain a final dataset of 11,290 high-quality gene models, 11,033 (98%) of which are placed within the 27 chromosome-scale scaffolds. This gene set is comparable in size to those annotated by the NCBI Eukaryotic Genome Annotation Pipeline for other ant genomes (range: 10,491-15,668; N= 18 different RefSeq ant genera; Table S1). We compared the obtained gene set of *C. hispanica* (Chis2) with 19 published ant annotations. Out of the 258,587 protein-coding genes analyzed using OrthoFinder (Emms and Kelly 2019), 96.82% (250,353) were placed in 13,698 orthogroups. Of these, 1,407 were species-specific and 6,199 were found in all species including 3,365 single-copy genes. The orthogroup profile of *C. hispanica* was overall comparable to that of other ants (Figure 4). However, our annotation had one of the smallest number of genes placed in orthogroups (10,918), and one of the largest proportions of unassigned genes (3.3%).

**Figure 4:**
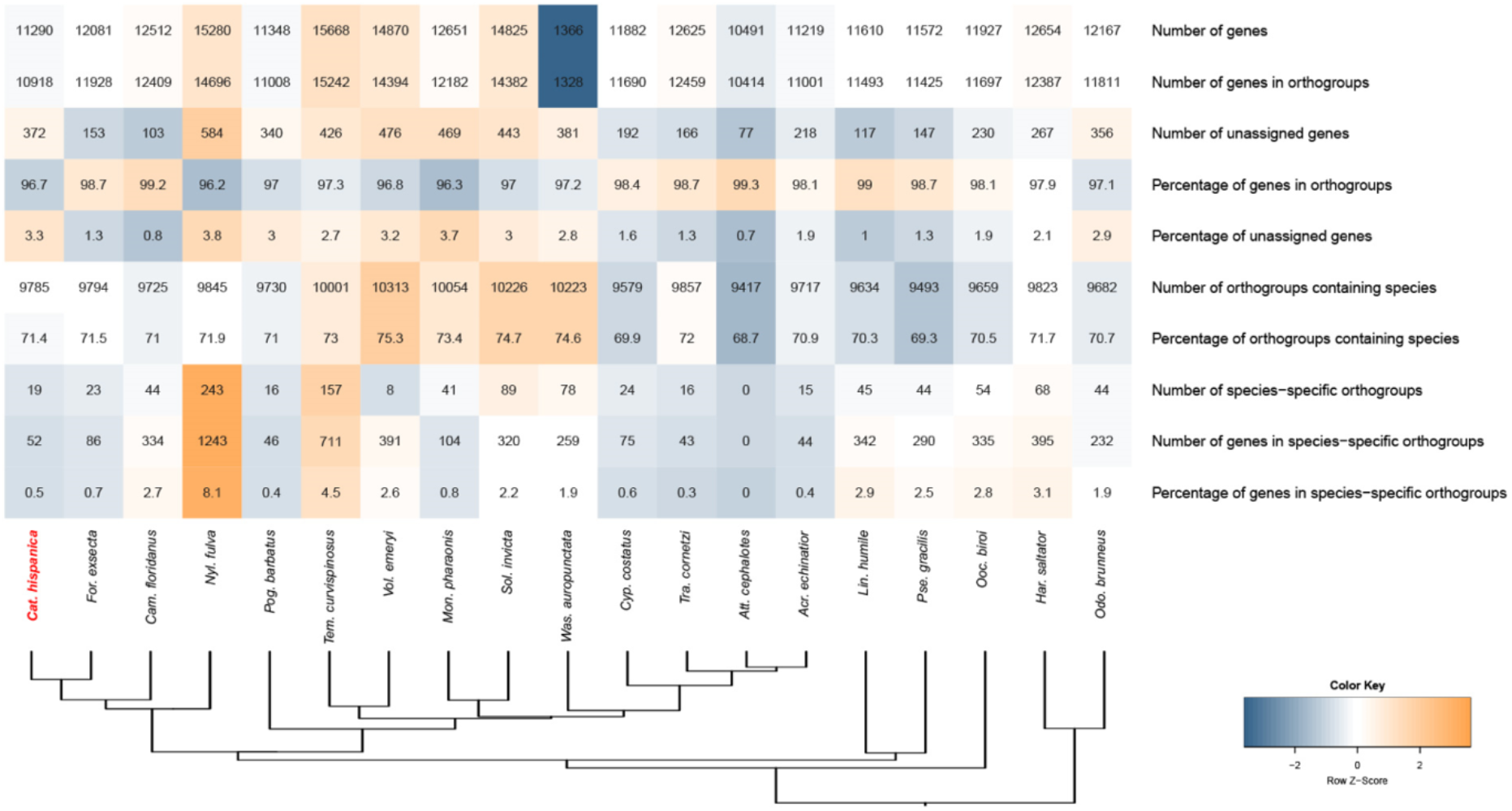
Summary values from the ortholog analyses. The color intensity indicates the z-score of variation (deviation from the mean) among all species, from the smallest value (blue) to the highest value (orange). Species are ordered according to their phylogenetic positions inferred from a concatenated alignment of single-copy orthologs.

### Repeat annotation

We built custom repeat libraries for each of the two assemblies of *C. hispanica* and for the 19 published ant genomes (see genome accessions in Table S1). The Chis1 and Chis2 assemblies contained 1,708 and 1,673 different repetitive elements, which accounted for 15.43% (31,851,170 bp) and 15.1% (31,512,815 bp) of their assembly sizes, respectively (Figure 5). A large proportion of these corresponded to unclassified interspersed repeats (6.7% / 6.78% of the genomes; Figure S6). The two genomes also contained 2.0% / 1.8% of Class I (retroelements), and 2.18% / 1.85% of Class II elements (DNA transposons). In total, 56 different families of repetitive elements were annotated in *C. hispanica*. LTR/Gypsy were the most frequent transposable elements of Class I in the genomes (0.53% / 0.82%), while large Polintons / Mavericks were the most abundant Class II transposable elements (0.98% / 0.67%). Across published ant assemblies, the total proportion of transposable elements appeared quite variable irrespective of their phylogenetic relationships (range: 17.27 – 48.47%; N= 19 ant species; Figure 5; Table S2). The *C. hispanica* assemblies had smaller proportions of repetitive elements (15.1% - 15.43%) than any of these assemblies, including that of *Formica exsecta* (18.53 %), the closest species available for comparison. The relatively low proportion of transposable elements observed in the genomes of *C. hispanica* may be due to the fact that it was assembled primarily from noisy nanopore long-reads, possibly leading to a collapse of repeated regions. Alternatively, *C. hispanica* may resist the invasion and proliferation of transposable elements more efficiently than other species. Whether its unusual reproductive system, combining both diploid and haploid parthenogenesis for queen and male production, could help keep transposable elements at bay deserves further exploration.

**Figure 5:**
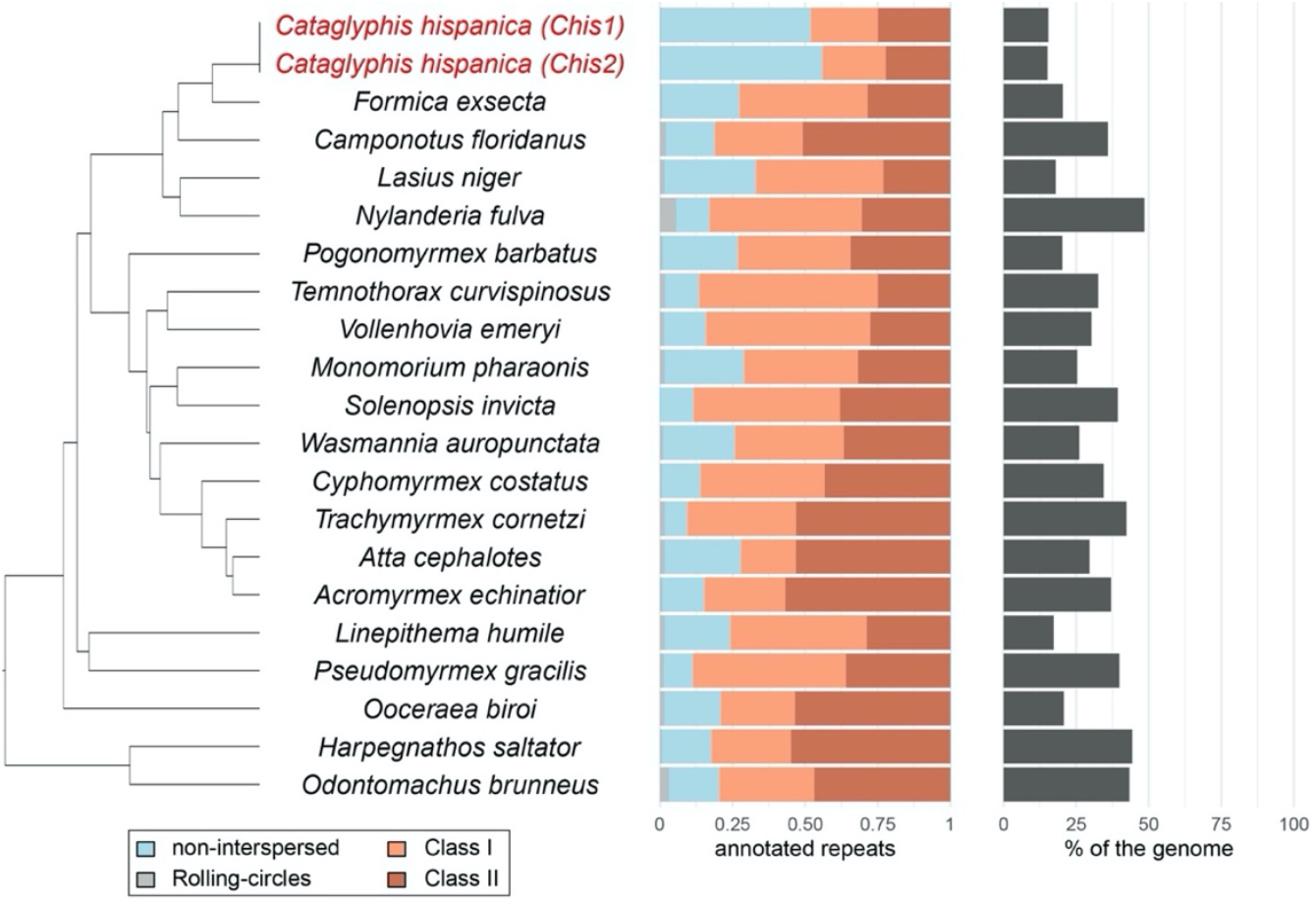
Summary of the repetitive elements’ categories annotated in 20 different ant species using our custom pipeline. The ratios of the major categories of repetitive elements identified in each species is shown on the left. The total proportion of repetitive elements found in each genome is shown on the right. Species are ordered accordingly to their phylogenetic positions inferred from a concatenated alignment of single-copy orthologs.

### Lineage comparison

We previously showed that *Cataglyphis hispanica* consists of two divergent lineages that are readily identifiable using microsatellite markers (Darras *et al*. 2014). Individuals from Chis1 and Chis2 lineages can however not be distinguished based on external traits: they share virtually the same morphologies for the queen and male castes, co-occur in the same localities and do not differ in any obvious colony characteristics. Furthermore, although queens only successfully produce workers when they mate with a partner originating from the other lineage, we have no evidence that lineages can recognize each other and avoid assortative mating. The interdependent nature of the lineages could stem from a small number of recessive mutations biasing development toward the queen caste in each lineage. Such “royal cheats” (Hughes and Boomsma 2008) seem common in eusocial Hymenoptera and have been hypothesized to be at the origin of caste determination and possibly social hybridogenesis (Anderson *et al*. 2008; Weyna *et al*. 2021; Withrow and Tarpy 2018). In line with their observed phenotypic similarity, assemblies of the two lineages appear highly similar and colinear (Figure S3). Large indel (>10kb) variation among lineages account for 6.6 % (13.6 Mb, Chis1) and 6.4 % (13.2 Mb, Chis2) of the chromosome-scale scaffolds. These “lineage-specific” indels are scattered across the assemblies (Figure S7) and are gene-deprived; only 35 of the 11,033 (0.3%) genes models from the Chis2 chromosome-scale scaffolds turned missing from Chis1 when performing an annotation lift-over using Liftoff (Table S3). Small inter-lineage polymorphism (i.e., SNPs and indels smaller than 100 bp) also appear uniformly distributed across chromosomes, with no large portion of chromosomes showing elevated divergence among assemblies (Figure S7). This later result contradicts previous hypotheses that hybridogenetic lineage pairs might be determined by ancient non-recombining regions, as found in other dimorphic system such as sex chromosomes or social chromosomes (Darras *et al*. 2014; Linksvayer *et al*. 2013; Schwander *et al*. 2014).

We additionally estimated divergence between the two genomes sequenced analyzing polymorphism at four-fold-degenerate sites, which are expected to be evolving neutrally since every mutation at a four-fold site is synonymous. Our annotation of the Chis2 genome contained 2,620,448 four-fold-degenerate sites. Among these, 13,048 had a different allele in the Chis1 and Chis2 males used to obtain haploid genome consensus. Assuming no recombination and a typical insect mutation rate of approximately 3 × 10^−9^ mutations per neutral site per haploid genome per generation (Keightley *et al*. 2014, 2015; Yang *et al*. 2015; Liu *et al*. 2017; Oppold and Pfenninger, 2017), this proportion of mutated four-fold-degenerate sites translated into an average divergence time of about 830,000 generations between the alleles of the two males sequenced (Obbard *et al*. 2012). Hence, the two genomes sequenced may have diverged almost 1 million years ago (assuming one generation per year) - a divergence time similar to that observed between closely related species of fire ants (Cohen and Privman, 2019). The origin of the hybridogenetic lineages themselves could be much younger though, considering they might have emerged from two divergent populations or shared ancestral polymorphism (Darras *et al*. 2019).

## Conclusion

We generated high-quality chromosome level genome assemblies of the two lineages of the hybridogenetic ant *C. hispanica*, a representative species of the thermophilic ant genus *Cataglyphis*. Using chromosome conformation capture, we identified a Robertsonian translocation between the two queens sequenced, resulting in 26 and 27 chromosomes, respectively. However, this difference in chromosome numbers seemed not fixed between lineages, suggesting that this chromosome rearrangement was not pivotal in the origin and maintenance of social hybridogenesis in *C. hispanica*. The two lineage assemblies were overall very similar with no large-scale region showing high divergence. Future work using population genomic approaches and genomic comparisons with other *Cataglyphis* species exhibiting social hybridogenesis will be necessary to identifying polymorphic genes or regulatory regions that are involved in the differentiation of queens and workers during development.

## Methods

### Biological samples

Permits were obtained to collect colonies of *Cataglyphis hispanica* in three Spanish locations (Bonares, Caceres and Merida; Figure 1B). Male samples from Bonares were used for Illumina DNA sequencing. Shortly after sampling, the Bonares population was wiped out by human activities. Consequently, samples from another locality (Caceres) were used for subsequent Nanopore sequencing, 3C-seq and RNA-seq. Male and worker pupae from two distant localities (Bonares and Merida) were used for karyotyping. Twelve diagnostic microsatellite loci were genotyped prior to sequencing and karyotyping to assess the lineage membership of each queen and male and to confirm that workers were all first generation hybrids (Darras *et al*. 2014).

### DNA and RNA-Sequencing

Genomic resources were generated for both the Chis1 and the Chis2 lineages. High-molecular-weight DNA was extracted from pure lineage queen and male individuals using QIAGEN Genomic-tips. For each lineage, two queen clones originating from the same nest were used for Nanopore sequencing. Queens of *C. hispanica* are produced through automictic parthenogenesis with central fusion which results into diploid individuals that are highly homozygous (Darras *et al*. 2014; Pearcy *et al*. 2011) and thus suitable for genome assembly. Nanopore libraries were prepared using rapid sequencing kits (SQK-RAD001 and SQK-RAD004). The resulting long read libraries were sequenced on MIN106 flow cells and basecalled using Albacore v2.1.10. For each lineage, three Illumina libraries were generated from whole-genome amplified DNA extracted from a single male with mean insert sizes of 170 bp, 500 bp and 800 bp, and sequenced with a HiSeq2000 (paired-end 2 × 100 bp mode).

3C-seq libraries were prepared according to the protocol described in Marie-Nelly *et al*. (2014). Briefly, queens from both lineages had their gut removed and were immediately suspended in 30 mL of formaldehyde solution (Sigma Aldrich; 3% final concentration in 1X tris-EDTA buffer). After one hour of incubation, quenching of the remaining formaldehyde was done by adding 10 mL of glycine (0.25 M final concentration) to the mix for during 20 min. The cross-linked tissues were pelleted and stored at −80°C until further use. The 3C-seq libraries were prepared using the *Dpn*II enzyme and sequenced using an Illumina NextSeq 500 apparatus (paired-end 2×75 bp; first ten bases corresponding to custom-made tags). 3C-seq libraries are similar to Hi-C libraries except that they contain a higher percentage of paired-end reads due to the lack of an enrichment step (Flot *et al*. 2015).

To help annotate the genomes, three normalized RNA-seq cDNA Illumina libraries were obtained: one from an adult Chis1 queen, one from a Chis2 queen and one from a brood pool comprising multiple developmental stages and adult workers originated from colonies of the two lineages (HiSeq2000, paired-end 2 × 100 bp mode).

### Genomes assembly

The genome of each hybridogenetic lineage was assembled independently following the pipeline depicted in Figure S1. Nanopore data were assembled using Flye v2.7 with four iterations of polishing based on long reads (Kolmogorov *et al*. 2019). Raw Illumina reads were trimmed for quality and adapters were removed using Trimmomatic v0.32 with options ILLUMINACLIP:TruSeq3-PE-2.fa:2:30:10 LEADING:3 TRAILING:3 SLIDINGWINDOW:4:15 MINLEN:36 (Bolger *et al*. 2014). The trimmed reads were then aligned to the long-read assemblies using BWA-MEM v0.7.15 (Li and Durbin 2009). SNPs and indels with at least three supporting observations were called using freebayes v1.2 (Garrison and Marth 2012), and error-corrected consensus sequences were obtained using BCFtools v1.4 (Li *et al*. 2009).

To obtain chromosome-scale assemblies, we scaffolded the polished contigs with the 3C reads using instaGRAAL, a MCMC, proximity-ligation based scaffolder (Baudry *et al*. 2020; Marie-Nelly *et al*. 2014). The 3C reads were trimmed using cutadapt (Martin 2011) and subsequently processed using hicstuff (Matthey-Doret *et al*. 2020) with the following parameters *–aligner bowtie2 –iterative –enzyme DpnII*. The instaGRAAL scaffolder was run on the pre-processed data for 100 cycles (parameters: level 4, with options *--coverage-std 1 –level 4 –cycles 100*) (Baudry *et al*. 2020) and final scaffolds were obtained using the instaGRAAL *-polish* script, with all corrective procedures at once (only one parameter: *-m polish*). Briefly, instaGRAAL explores the chromosome structures by testing the relative positions and/or orientations of DNA segments (or bins) according to the contacts expected given a simple three-parameter power-law model. These modifications take the form of a fixed set of operations (swapping, flipping, inserting, merging, etc.) of bins corresponding to 3^4^ = 81 *Dpn*II restriction fragments. The likelihood of the model is then maximized by sampling the parameters using a MCMC approach (Marie-Nelly *et al*. 2014). After 100 iterations (i.e., a likely position for each bin is tested 100 times), the genome structure converges towards a relatively stable structure that does not evolve anymore when more iterations are added, resulting in chromosome-level scaffolds. The algorithm is probabilistic and ignores initially part of the intrinsic structure of the original contigs in order to sample a larger range of genome space (Baudry *et al*. 2020). Therefore, some trustworthy information contained in the initial polished assembly can be lost, or modified, along the way. The final correction step of instaGRAAL consists in reintegrating this lost information into the final assembly, to correct for instance local untrustworthy tiny inversions of individual bins within a contig. The contact maps of the scaffolded assemblies were built using hicstuff. Gaps created during the scaffolding process were closed using Nanopore data with four iterations of TGS-GapCloser (Xu *et al*. 2019) and new polished consensus sequences were obtained using BCFtools (see method above). Completeness of the assemblies were assessed at each step using BUSCO v5.2.2 with the Hymenoptera odb10 lineage (Simão *et al*. 2015; Waterhouse *et al*. 2017). We also ran KAT v2.4.1 to compare the k-mer frequencies of Illumina reads to final assemblies (Mapleson *et al*. 2016). To investigate differences in chromosomal arrangement among lineages, the two genome assemblies were aligned with minimap2 v2.17 (exact preset: -x asm5) and alignments were visualized using dot plots obtained with D-GENIES (Cabanettes and Klopp, 2018).

### Karyotyping

To validate the number of chromosomes inferred from 3C contact information, chromosome preparations were made from brains of male and worker larvae following the protocol described by (Lorite *et al*. 1996), with some modifications. Briefly, larvae at the last instar stage were dissected and their cerebral ganglia were transferred to microplate wells with 0.05% colchicine in distilled water. After 30 min, samples were transferred to a fixative solution (acetic acid:ethanol, 3:1) and incubated for 45 min. Ganglia cells were disaggregated in a drop of 50% acetic acid on a clean slide, new fixative solution was added, and the slides were dried at 60ºC. Chromosome preparations were stained with 10% Giemsa in phosphate buffer (pH 7). Microscopy images were captured with a CCD camera (Olympus DP70) coupled to a microscope (Olympus BX51) and were processed using Adobe Photoshop.

### Gene annotation

We used the Chis2 chromosome-level assembly for gene annotation. A repeat library was constructed using the REPET package v2.5 (Flutre *et al*. 2011; Quesneville *et al*. 2005). This repeat library was cleaned up manually to remove bacterial genes, mitochondrial genes and genes with hits to the gene set of the ant *Cardiocondyla obscurior* (v1.4) which had been purged of transposable elements (Schrader *et al*. 2014). The fraction of the genome classified by RepeatClassifier as “Unknown” was reduced from 2.2% to 0.9% as a result of this procedure. Repeats were soft-masked using RepeatMasker v4.0.7 (Smit and Hubley, http://www.repeatmasker.org) prior to *de novo* gene prediction.

Gene models were inferred from RNA-seq, homology data and *ab initio* predictions. The three RNA-seq libraries were aligned to the *Chis2* genome using STAR v2.6.0 (Dobin *et al*. 2013) with the multi-sample 2-pass mapping strategy. Transcripts were then assembled using Trinity v2.10.0 (Grabherr *et al*. 2011; Haas *et al*. 2013)(options --genome_guided_max_intron 100000 --jaccard_clip) and combined into gene models using PASA (Haas *et al*. 2003). Ant proteomes annotated using the NCBI Eukaryotic Genome Annotation pipeline (RefSeq, taxid:36668) were aligned to the genome using GenomeThreader v1.5.10 (Gremme *et al*. 2005) in order to predict gene structures. AUGUSTUS *ab initio* predictions were generated using BRAKER v2.1.02 (Hoff *et al*. 2016, 2019) based on hints from RNA-seq data and GenomeThreader protein alignments (--etpmode). BRAKER was first run with preliminary AUGUSTUS parameters trained by running BUSCO v3.0.2 on the genome assembly (--long option; Hymenoptera odb9 database). To refine the training of AUGUSTUS, the most accurate gene models inferred by BRAKER were then identified using GeneValidator (Drăgan *et al*. 2016) with RefSeq ant proteomes as references and an arbitrary quality threshold of Q89. To avoid biases, predicted proteins with more than 70% sequence identity to another protein in the set were removed from the selected gene models using the aa2nonred.pl script provided with BRAKER. The resulting gene models were used to train AUGUSTUS again, and BRAKER was run with the new parameter set. *Ab initio*, RNA-seq-based and homology-based gene predictions were combined into a single gene set using EvidenceModeler v1.1.1 (Haas *et al*. 2008) with the following weight settings: PASA alignments: 10; GenomeThreader alignments: 3, Augustus predictions: 1, PASA/Transdecoder predictions: 1, GenomeThreader predictions: 1. Functional information was obtained from eggNOG-mapper v2 (Huerta-Cepas *et al*. 2017, 2019) with the options “taxonomic scope adjusted per query” and “annotations transferred from any ortholog”. Protein sequences with similarity to RefSeq ant proteins (as of July 2019) were identified using blastp and an E-value threshold of 10^−5^. Annotations with no known functional information and no hits to any RefSeq ant proteins were filtered out.

### Comparative analyses

To identify orthologous and taxonomically restricted genes, we compared the proteomes of *C. hispanica*, of 18 ants annotated by the NCBI Eukaryotic Genome Annotation Pipeline (Table S1) and of *Lasius niger* (Konorov *et al*. 2017) using OrthoFinder v2.3.12 (Emms and Kelly, 2019) with its standard DEndroBLAST workflow. We used the feature annotation tables from RefSeq annotations to select the longest isoform of each gene annotated by NCBI prior to analysis. The published genome of *L. niger* is highly incomplete (no more than 65% of the 4,415 highly conserved single-copy genes of BUSCO’s Hymenoptera odb9 database are found in this assembly). Consequently, it was only used to guide phylogenetic analyses due to its relative proximity with *Cataglyphis*. A preliminary catalog of single-copy orthologs was obtained from a first run of OrthoFinder. Single-copy sequences were aligned with Mafft v7.310 (Katoh and Standley, 2013) and the alignments were trimmed with trimAL v1.4.1(options “-gt 0.8 -st 0.001”) (Capella-Gutiérrez *et al*. 2009). The concatenated alignments were then passed to IQ-TREE v1.7.17 (option “-m LG+R4”) (Nguyen et al. 2015) to infer a species tree. The tree was converted to an ultrametric topology with the r8s program with options “mrca root Obir Hsal; fixage taxon=root age=150; divtime method=LF algorithm=TN” (Sanderson 2003). The resulting species tree was used for a second, more precise run of OrthoFinder.

### Repeat annotation

To compare the frequency of repetitive elements found in the genome of *C. hispanica* to the frequencies found in the genomes of other ant species available (Table S2), we constructed optimized repeat libraries for each species using a custom pipeline (https://github.com/nat2bee/repetitive_elements_pipeline). Shortly, repeat libraries were built with RepeatModeler v1.0.11 (http://www.repeatmasker.org/RepeatModeler/), TransposonPSI (http://transposonpsi.sourceforge.net/) and LTRharvest from GenomeTools v1.6.1 (Ellinghaus *et al*. 2008). For each species, the different libraries were merged into a non-redundant library (<80% identity) using USEARCH v11.0.667 (Edgar 2010). Library annotations were obtained with RepeatClassifier. Each custom library was concatenated with the Dfam v3.1 Hymenoptera library of RepeatMasker v4.1.0 and used to annotate repeats in the genome of the corresponding species using RepeatMasker. Summary statistics of the annotated repeats were obtained with RepeatMasker_stats.py (https://github.com/nat2bee/repetitive_elements_pipeline).

### Lineage comparison

The two assemblies were aligned with minimap v2.19 (-cx asm5 –cs) and variants were called with paftools (paftools.js call -L5000 -l1000). The distribution of large indels (>10 kb) and the density of small polymorphisms (SNPs and indels no larger than 100 bp) across the genomes were calculated using custom scripts. Annotation lift-over from the Chis2 assembly on to the Chis1 assembly was performed with Liftoff v1.6.3 (Shumate and Salzberg 2020). To verify if missing annotations did not result from misassemblies, we also lift these on a consensus Chis1 assembly derived from alignment of the Chis1 haploid short reads on the Chis2 assemblies using BCFtools as described above (see Genomes assembly) with regions not covered by reads masked to avoid reference bias (--mask --mask-with N).

To estimate the divergences of the two lineages of *C. hispanica*, we investigated the polymorphism at 4-fold-degenerate sites, which we assumed to be neutrally evolving. The Illumina read of the Chis1 lineage were mapped onto the Chis2 reference genome and single-nucleotide variants were called using MapCaller v0.9.9.41 (Lin and Hsu 2019). The resulting vcf file was filtered to keep only single-nucleotide variants with two alleles and a ‘PASS’ quality filter. To determine the proportion of 4-fold sites that were polymorphic among our male samples of the two lineages, the positions of 4-fold sites in coding sequences of our annotation were identified using a custom script (T. Sackton, https://github.com/tsackton/linked-selection).

## Acknowledgements

We thank A. Cohanim, E. Privman and R. Faure for their advice on early genome assemblies, F. Rodriguez and L. Grumiau for their assistance with genotyping and Q. M. Pan for her comments on the manuscript. Version 3 of this preprint has been peer-reviewed and recommended by *Peer Community In Genomics* (https://doi.org/10.24072/pci.genomics.100017).

## Data, scripts and codes availability

All the raw sequencing data and genome assemblies generated during this study have been deposited at NCBI (Accession numbers: SRR17481978 – SRR17481992). The genomes of *C. hispanica* were deposited in NCBI (Accession numbers: JAJUXC000000000 and JAJUXE000000000). Codes are available at figshare (https://doi.org/10.6084/m9.figshare.17964695.v7).

## Supplementary material

Supplementary figures, tables, gene annotations, TE repeat libraries and reports can be accessed at figshare (https://doi.org/10.6084/m9.figshare.17964695.v7).

## Authors’ Contributions

HD collected the ants, prepared DNA/RNA, performed Nanopore sequencing (together with JFF), assembled and annotated the genome. NSA performed TE analyses and genomic comparisons. Both HD and NSA prepared first manuscript draft. PL performed karyotyping. LB optimized 3C scaffolding parameters. NG performed 3C scaffolding. MM prepared the 3C libraries. FR constructed the TE library. IA supervised the construction of the TE library. RK supervised 3C library generation and scaffolding. JFF participated in the Nanopore sequencing and supervised genome assembly. SA collected the ants, designed, and supervised the study. All authors read and approved the final manuscript.

